# Massive bursts of transposable element activity in Drosophila

**DOI:** 10.1101/010231

**Authors:** Robert Kofler, Viola Nolte, Christian Schlötterer

## Abstract

The evolutionary dynamics of transposable element (TE) insertions have been of continued interest since TE activity has important implications for genome evolution and adaptation. Here, we infer the transposition dynamics of TEs by comparing their abundance in natural *D. melanogaster* and *D. simulans* populations. Sequencing pools of more than 550 South African flies to at least 320-fold coverage, we determined the genome wide TE insertion frequencies in both species. We show that 46 (49%) TE families in *D. melanogaster* and 44 (47%) in *D. simulans* experienced a recent burst of activity. The bursts of activity affected different TE families in the two species. While in *D. melanogaster* retrotransposons predominated, DNA transposons showed higher activity levels in *D. simulans*. We propose that the observed TE dynamics are the outcome of the demographic history of the two species, with habitat expansion triggering a period of rapid evolution.

## Introduction

The question of how the dynamics of transposable elements (TEs) develop over evolutionary time scales has not yet been resolved in a satisfactory manner. Rather, two competing models are proposed: The equilibrium model assumes that genomes are in transposition-selection balance and TEs have a constant transposition rate. With negative selection counteracting the spread of transposable elements (Charlesworth and Langley, 1989; Petrov et al., 2003; Lockton et al., 2008; Petrov et al., 2011; González et al., 2009; Lee and Langley, 2010) the TE composition remains stable over time. By contrast, the non-equilibrium model rests on a variable TE activity over evolutionary time scales (Kofler et al., 2012; Blumenstiel et al., 2013; Le Rouzic et al., 2007; Bergman and Bensasson, 2007; Lerat et al., 2011; El Baidouri and Panaud, 2013). Consistent with this model, bursts of activity have been shown for some TE families (Choulet et al., 2010; Engels, 1992; Diez et al., 2014; Bailey et al., 2003). It is, however, not clear whether bursts are limited to a few TE families within a species, or if they are a general phenomenon of all, or at least the majority of TE families. The controversy about the dynamics of TEs has been particularly hard to resolve from single genomic sequences or population data of a single species, because both models, the equilibrium and the non-equilibrium model, lead to similar predictions. The negative correlation between the number of insertions of a TE family and the average population frequency provides a good example for this (Petrov et al., 2003; Kofler et al., 2012). Under the equilibrium model the low average population frequency of abundant TE families is caused by strong purifying selection (Petrov et al., 2003, 2011). The same observation, however, can be attributed to a recent burst of TE activity under the non-equilibrium model (Kofler et al., 2012; Bergman and Bensasson, 2007). While the pattern of genome-wide TE insertions and their frequency distribution in a single species are not sufficient to resolve this controversy, a comparison of the TE abundance in two closely related species can do so, since the two models lead to different predictions: assuming a constant TE activity since the split of two closely related species, as purported by the equilibrium model, a very similar TE abundance should result in both species (Petrov et al., 2003). Under a non-equilibrium model bursts of TE activity are independent after the species split, resulting in divergence in TE insertion patterns between the two species. In this study we investigated the TE content in natural *D. melanogaster* and *D. simulans* populations, two closely related species which diverged about 2-3 million years ago (Lachaise et al., 1988; Hey and Kliman, 1993). Combining empirical TE insertion frequency estimates from Pool-Seq (Schlötterer et al., 2014) with computer simulations of TE dynamics we identify for about 78 (84%) TE families a significant deviation from the equilibrium model. About 46 (49%) TE families in *D. melanogaster* and 44 (47%) families in *D. simulans* experienced a recent burst of activity. Interestingly, retrotransposon families had a higher rate of transposition bursts in *D. melanogaster* while DNA transposons were more active in *D. simulans*. We propose that the high rate of non-equilibrium dynamics may be the result of the recent colonization of these two species.

## Results

We compared the TE abundance in natural populations of the two closely related species *D. melanogaster* and *D. simulans* to determine the patterns of long-term transposition rates. In particular, we used these data to distinguish between a continuous rate of transposition and variable rates, with periods of burst followed by low activity. The comparison of TE abundance in the two species has been complicated by markedly different qualities of the reference genomes and the associated TE annotations. To avoid that the higher quality of the *D. melanogaster* genome biases our results, we pursued the following strategies: (i) using an improved *D. simulans* reference assembly (Palmieri et al., 2014), (ii) restricting the TE abundance comparison to regions present in the assemblies of both species (iii) using the same *de novo* TE annotation pipeline in both species (see Material and Methods) (iv) and employing a TE calling method, which is independent of the presence of a TE insertion in the reference genome. From each species we analyzed isofemale lines collected 2013 in Kanonkop (South Africa). By sequencing pooled individuals (Pool-Seq) (Schlötterer et al., 2014) we obtained an average coverage of at least 320-fold using Illumina paired end reads. We estimated TE abundance from the corresponding average physical coverage of 145 at TE insertion sites, using PoPoolation TE (Kofler et al., 2012).

A comparison of *de novo* annotated TEs in *D. melanogaster* with the reference annotation [FlyBase; v5.53; (Quesneville et al., 2005; Kaminker et al., 2002)], indicated that our pipeline has a high sensitivity as well as a high specificity (supplementary results 3.1). The high quality of our TE annotation is further supported by similar population frequency estimates (Spearman’s rank correlation, *r*_*S*_ = 0.82, *p* < 2.2*e* − 16) and insertion numbers (Spearman’s rank correlation, *r*_*S*_ = 0.81, *p* < 2.2*e* − 16; supplementary results 3.3) in *D. melanogaster* , between this and a previous study that used the reference annotation (Kofler et al., 2012). As final validation of our annotation pipeline we compared the genomic TE distribution in natural populations obtained from our pipeline to an independently acquired data set. Vieira *et al.* (1999) estimated the abundance of 36 TE families in *D. melanogaster* and *D. simulans* populations by *in situ* hybridization. We obtained a high correlation between the estimates of both methods (*D. melanogaster* : Spearman’s rank correlation, *r*_*S*_ = 0.85, *p* = 3.6*e* − 9; *D. simulans*: *r*_*S*_ = 0.62, *p* = 0.0002; supplementary results 3.3), confirming the robustness of our method.

The number of TE insertions differs markedly between the two species (fig. 1; supplementary table 5), with a larger number of TE insertions in *D. melanogaster* (18, 382) than in *D. simulans* (13, 754). A similar observation has been made previously using *in situ* hybridization data (Dowsett and Young, 1982; Aquadro et al., 1988; Vieira et al., 1999). The average population frequency of TE insertions is higher in *D. simulans* (0.199) than in *D. melanogaster* (0.146), mostly due to a higher number of TE insertions with intermediate to high frequency in *D. simulans* (> 0.2; supplementary fig. 1). Analyzing the different TE classes separately we uncovered pronounced differences between the two species. *D. melanogaster* has markedly more Long Terminal Repeat (LTR; *D.mel.* = 7,252, *D.sim.* = 3,222) and non-LTR (*D.mel.* = 5,723, *D.sim.* = 2,902) insertions, whereas *D. simulans* has more Terminal Inverted Repeat (TIR) insertions (*D.mel.* = 5,021, *D.sim.* = 7,258). Many RNA transposon families (LTR and non-LTR) have more insertions in *D. melanogaster* whereas DNA transposon families (TIR) are more frequently inserted in *D. simulans* (fig. 2). The unexpected presence of the P-element in *D. simulans* [fig. 2; (Brookfield et al., 1982; Engels, 1992; Vieira et al., 1999)] will be discussed elsewhere (Kofler *et al.*; in preparation).

To answer whether the equilibrium or non-equilibrium dynamics predominated in the evolution of TEs of *D. simulans* and *D. melanogaster* we require two pieces of information, first the age distribution of TE insertions and second, whether differences in insertion numbers between the two species significantly deviate from equilibrium expectations.

**Figure 1:**
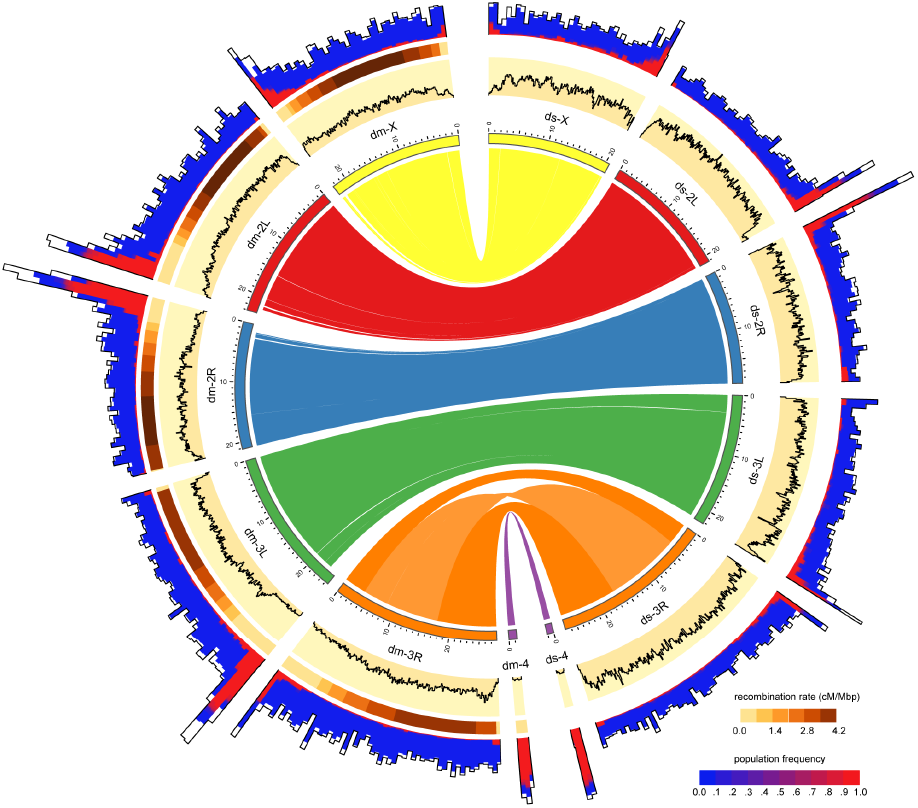
Distribution of TE insertions in a natural population of *D. melanogaster* (dm) and of *D. simulans* (ds). The TE distribution (outer graph) is compared to the recombination rate (middle graph) and the nucleotide polymorphism (Θ_*π*_, yellow inner graph). TE abundance and recombination rate are shown for windows of 500kb, whereas the nucleotide diversity is shown for windows of 100kb. For overlapping TE insertions (white) no estimates of population frequencies could be obtained. The relationship between the reference genomes is shown in the inside. Note, the inversion on chromosome 3R (Sturtevant, 1921) and the missing pericentromeric regions in the assembly of *D. simulans*. The maximum nucleotide diversity of the plot is 0.018 and the maximum number of TE insertions 400

**Figure 2:**
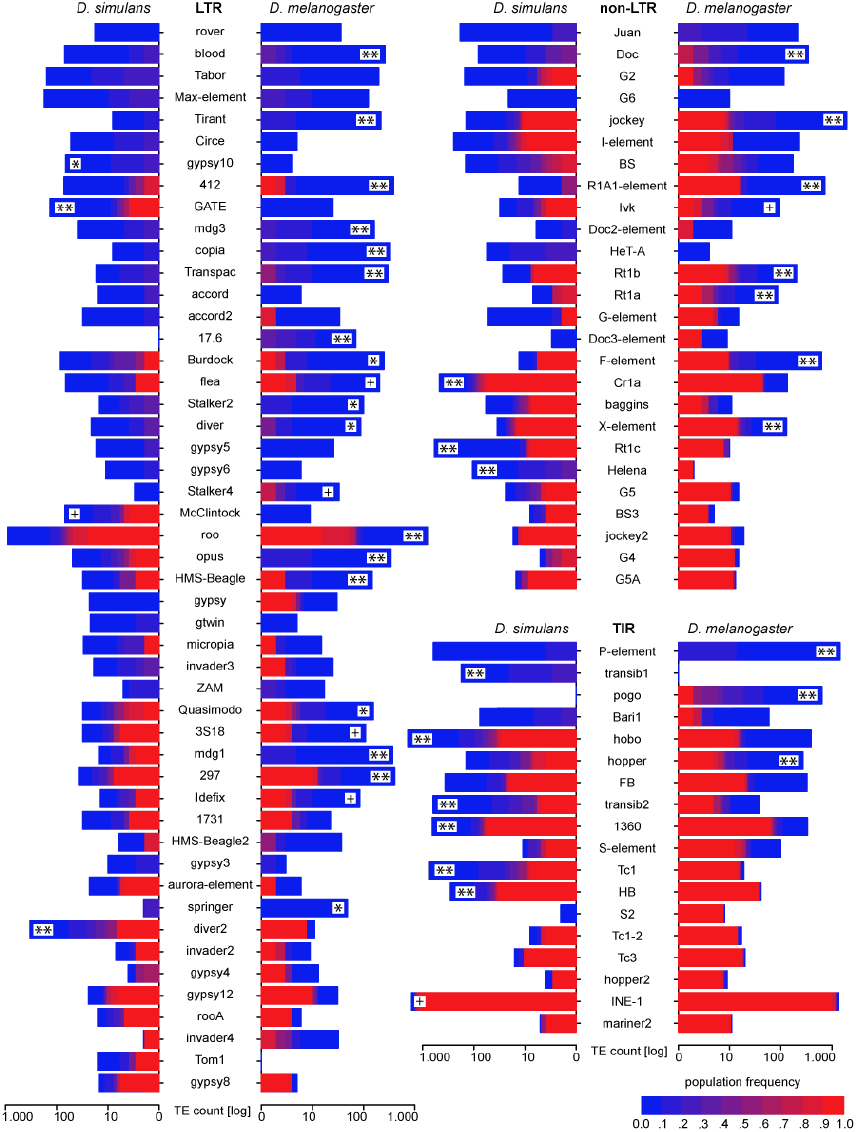
Abundance of different TE families in a natural *D. melanogaster* and of *D. simulans* populations; Significant deviation of TE insertion numbers from expectations under an equilibrium model is indicated for the species with the higher number of insertions. p-values after Bonferroni correction: ** < 0.001; * < 0.01; + < 0.05; Only TE families with more than 10 insertions are shown. Foldback (FB) is grouped with TIRs solely for graphic reasons

The age distribution is an important parameter describing the dynamics of transposable elements. A direct approach to determine the age of TE insertions is based on the number of mutations after insertion (Bergman and Bensasson, 2007; Malik et al., 1999; Blumenstiel et al., 2002, 2013), but this method cannot be applied to Pool-Seq data. Nevertheless, the previously demonstrated strong correlation between sequence divergence of TEs and their frequency in a natural population (Kofler et al., 2012) suggests that population frequencies of TE insertions are good age estimators: young insertions have a low population frequency, while old insertions tend to be fixed. We further scrutinized this relationship by reasoning that young TE insertions are more likely to be expressed. Using RNA-Seq data from *D. simulans* (Palmieri et al., 2014) we found a highly significant negative correlation (Spearman’s rank correlation, *r*_*S*_ = *−*0.34, *p* = 0.00024) between population frequency and expression intensity. Finally, we reasoned that fixed TE insertions are old and therefore more likely to be shared between species. Indeed, we found fixed TE insertions to be highly enriched for TE insertions shared between *D. melanogaster* and *D. simulans* (Fisher’s exact test; *p* < 2.2*e* − 16; supplementary results 3.6). Thus these fixed TE insertions most likely predate the split between these two species about 2-3 million years ago (Lachaise et al., 1988; Hey and Kliman, 1993). Overall, our analyses suggested that the population frequency of TE insertions provides a rough, but suitable estimator for the age of TE insertions.

We performed computer simulations to determine for each TE family whether the observed interspecific differences in TE insertions numbers between species (fig. 2) significantly deviate from expectations under an equilibrium model. Our simulations considered each TE family separately and relied on a fitness function in which fitness decreases exponentially with insertion number, a condition for obtaining stable equilibria (Charlesworth and Charlesworth, 1983). Given the strong influence of population size on TE dynamics (Lockton et al., 2008; Lynch and Conery, 2003) (supplementary fig. 2C), we used a population size ratio in our computer simulations that reflects the ratio of the population variation estimator *π* (*π^Dsim^/π^Dmel^* = 0.0113/0.0074 = 1.519; supplementary results 3.5). In about 50% (46/93) of the TE families the number of insertions deviated significantly from the expectations under an equilibrium model (fig. 2). This result was robust with respect to the actual population size employed in the computer simulations (for *N*_*e*_ > 10,000; supplementary results 3.9). Also when assuming an equal population size of the two species (e.g. (Nolte and Schlötterer, 2008)) substantial deviations from the equilibrium model were identified (supplementary fig. 3).

Based on the age distribution and the significance of the difference in insertion number between species, we addressed the question whether equilibrium or non-equilibrium dynamics of TEs predominate in the two species. TE families with a constant activity since the split of *D. melanogaster* and *D. simulans*, as purported by the equilibrium model, should have i) insertion numbers between the two species that do not significantly deviate from expectations under an equilibrium model, ii) both old and young insertions in both species and iii) roughly equal amounts of old insertions between the two species and iv) roughly equal amounts of young insertions between the two species. Only 15 (16%) of the TE families showed a TE distribution compatible with the equilibrium model (e.g.: I-element, BS; supplementary results 3.7; fig. 1). The remaining 78 (84%) families did not fit the expectations of the equilibrium model: 36 of them became only recently active (e.g.: rover, Juan), 5 were active only in one species (e.g.: Tom1, pogo), 12 were active only in the past (e.g.: invader4, BS3, INE-1) and 25 families had insertion numbers that differed significantly between the two species (e.g.: Burdock, jockey, hobo; supplementary results 3.7; fig. 1).

The large number of species specific TE activity patterns encouraged us to evaluate the distribution of TEs between two *D. melanogaster* populations from South Africa and Portugal. Consistent with the non-equilibrium model we observed substantial differences in TE abundance for two families (R1A1-element, gypsy2; supplementary results 3.3). This pattern is in agreement with previous observations (Biémont et al., 2003; Vieira et al., 1999) suggesting that the TE composition of local Drosophila populations can differ markedly despite little differentiation among cosmopolitan *D. melanogaster* populations (Caracristi and Schlötterer, 2003).

Given this high incidence of bursts of TE activity, we used the frequency based age estimates to distinguish between bursts of novel TE families (probably introduced by horizontal transfer (Sanchez-Gracia et al., 2005; Bartolomé et al., 2009)); no old insertions) and reactivation of old TE insertions. Reactivated families have old insertions in both species and the insertion numbers in the two species deviate significantly from equilibrium expectation (mostly due to differences in young insertions). We note that a highly active family with a rapid decrease in activity in one species may result in the same signature, but we consider this less likely since it is unclear whether high TE activity can be maintained for extended periods of time (Marí-Ordóñez et al., 2013; Khurana et al., 2011). We identified 28 bursts of novel families in *D. melanogaster* and 35 in *D. simulans*. 18 bursts of reactivated families could be detected in *D. melanogaster* and 9 in *D. simulans* (supplementary results 3.8). In total, 46 (49%) TE families in *D. melanogaster* and 44 (47%) in *D. simulans* experienced a recent burst of activity. Interestingly, while the amount of bursts of novel families is fairly equally distributed among the two species (*D.mel.*: LTR 24, non-LTR 3; TIR 1; *D.sim.*: LTR 23, non-LTR 8, TIR 4), *D. melanogaster* has more bursts of reactivated retrotransposon families while *D. simulans* has more bursts of reactivated DNA transposon families (*D.mel.*: LTR 9, non-LTR: 8, TIR 1; *D.sim.*: LTR 1, non-LTR 2, TIR 6; Fisher’s exact test: *p* = 0.0017).

## Discussion

In this report, we provide the first genome-wide characterization of TE abundance in large population samples of the two closely related species *D. simulans* and *D. melanogaster* . Consistent with previous reports (Vieira et al., 1999; Lerat et al., 2011), we found considerable differences in TE composition between the two species.

The new high quality TE annotation in *D. simulans* enabled us to address the longstanding debate about the TE dynamics in natural populations: does the genomic distribution of TEs represent a (stable) transposition-selection equilibrium or is the TE composition changing over time due to bursts of transposition activities (non-equilibrium model) (Charlesworth and Langley, 1989; Petrov et al., 2003; Lockton et al., 2008; Petrov et al., 2011; González et al., 2009; Lee and Langley, 2010; Kofler et al., 2012; Blumenstiel et al., 2013; Le Rouzic et al., 2007; Bergman and Bensasson, 2007; Lerat et al., 2011; El Baidouri and Panaud, 2013). With genome-wide TE insertion frequency data from populations of two species, we were able to distinguish these two hypotheses. Only 15 out of 93 TE families (16%) showed a pattern of TE abundance for which the equilibrium model could not be rejected. This is probably a conservative estimate since some episodes of TE burst may not be identified in our tests. For example, the distribution of I-element insertions seems to fit the equilibrium model, but it was suggested that active copies of the I-element were lost in *D. melanogaster* already some time ago, and that active copies only recently reinvaded extant populations (Bucheton et al., 1992). We note however, that another study did not find evidence for horizontal transfer of the I-element (Sánchez-Gracia et al., 2005).

Our strong support for a non-equilibrium model with massive bursts of TE activity depends to a large extent on our ability to date TE insertions correctly. In absence of sequence divergence data we relied on population frequency as an indicator of age. In addition to several lines of evidence supporting this approximation, it is remarkable that our results agree very well with the published literature: INE-1, jockey2, helena, Cr1a (but not baggins) were mainly active in the distant past (> 3 mya) in *D. melanogaster* (Kapitonov and Jurka, 2003; Singh and Petrov, 2004; Bergman and Bensasson, 2007), while 17.6, Stalker4, rover, flea, copia, mdg3, roo, Transpac, opus, blood, 412, Burdock, diver, Tirant, Juan, Doc were mostly active recently (< 100,000 ya) (Bergman and Bensasson, 2007; Engels, 1992). An example for some conflict with published evidence is the ZAM element, which was previously classified as an ancient element in *D. melanogaster* (Baldrich et al., 1997) but we identified only young insertions.

Interestingly, differences in TE composition are not only recognized in between species comparisons, but can be also detected between two *D. melanogaster* populations (supplementary results 3.3). These differences can not be explained by demographic events alone which should affect all TE families equally, whereas we only found marked differences for two TE families. Such differences in TE abundance between populations have also been observed in *D. simulans* (Biémont et al., 2003). In combination with the compelling evidence for bursts of TE activities in other species (Choulet et al., 2010; Engels, 1992; Diez et al., 2014; Bailey et al., 2003), we conclude that the TE composition in *D. simulans* and *D. melanogaster* is probably highly dynamic and changes quickly, such that even differences between populations can be detected. In fact at least 46 (49%) TE families in *D. melanogaster* and 44 (47%) in *D. simulans* have experienced a recent burst of transposition activity.

It is not clear what triggers bursts of TE activity. One interesting hypothesis suggests that habitat expansions could induce TE bursts (Vieira et al., 1999; Vieira and Biémont, 2004; Vieira et al., 2012). With TE insertions frequently contributing to adaptation to novel environments (Casacuberta and González, 2013; Kofler et al., 2012; González et al., 2008), TE bursts may be an important component of successful habitat expansions. Colonization of new environments may trigger bursts of TE activity by two, not mutually exclusive mechanisms. Either stress associated with new environments disturbs the guarding system, such as piRNA, or the habitat expansion may bring species into contact, which never met before. In combination with horizontal transfer of TEs, this could result in activity of a TE in a new host (Engels, 1992; Plasterk et al., 1999). One classic example for this scenario is the transfer of the P-element from *D. willistoni* to *D. melanogaster*, which invaded the territory of *D. willistonii* in South America (Engels, 1992). After the horizontal transfer, the P-element rapidly spread in *D. melanogaster* populations worldwide (Anxolabéhère et al., 1988). However, upon the activation of a single TE family, previously dormant families may become reactivated, as it has been noted during hybrid dysgenesis (Khurana et al., 2011; Petrov et al., 1995), where DNA damage mediated stress seems to be causative (Petrov et al., 1995; McClintock, 1984; Khurana et al., 2011). While we cannot pinpoint the most important mechanism for the burst of TE insertions, our data clearly indicate the central role of horizontal transfer. We identified transposition bursts of several novel TE families (28 in *D. melanogaster* and 35 in *D. simulans*), which are unlikely to be the outcome of de novo birth of TEs. Our results are fully compatible with previous work emphasizing the importance of horizontal transfer of TEs for the evolution of eukaryotic genomes (Sánchez-Gracia et al., 2005; Schaack et al., 2010; Gilbert et al., 2010). One key assumption for the habitat expansion mediated activity burst scenario is that the South African population does not represent an ancestral African population. In *D. melanogaster* , South African populations have been described to have high similarity to cosmopolitan ones (Pool and Aquadro, 2006). We also note that the level of polymorphism in our South African *D. simulans* population is more similar to a Portuguese population (*F*_*ST*_ = 0.030; a detailed analysis will be published elsewhere) than to a central African population (*F*_*ST*_ = 0.055; data from (Nolte and Schlötterer, 2008)). However, even very limited admixture from cosmopolitan flies would probably have been sufficient to introduce active TE families, resulting in a burst of TE activity.

The central role of habitat expansions for TE bursts, raises the question of the genomic TE distributions in species that remained in their original habitat. Does this imply that their genomic distribution resembles the predictions of an equilibrium model or does the immigration of non-native species also affect the TE distribution in native species? The analysis of ancestral African *D. melanogaster* and *D. simulans* populations may shed some light on this question as well as *D. sechellia* and *D. mauritiana*, two species that remained in their ancestral habitat. Furthermore, long read sequencing could provide a better characterization of recently inserted TEs (McCoy et al., 2014), which facilitates the timing of the transposition event.

Despite the rather frequent occurrence of horizontal transfer, several TE families experiencing a recent burst (i.e.: recent transfer) are restricted to one of the two species in our study (Tom1, 17.6, transib1, pogo; fig. 2). Some insights about this asymmetry come from another interesting difference between *D. simulans* and *D. melanogaster* . While *D. melanogaster* experienced more bursts of retrotransposon families, in *D. simulans* bursts of DNA transposons predominate. This apparent contrast could be the outcome of a different propensity for horizontal transfer among the major TE groups (LTR, non-LTR, TIR) in combination with the different colonization times of *D. melanogaster* and *D. simulans*. DNA transposons (TIR) and LTR transposons seem to be more prone to horizontal transfer than non-LTR TEs, since their double stranded DNA intermediates may be more stable than the RNA intermediate of non-LTR TEs (Schaack et al., 2010; Malik et al., 1999). Furthermore, the integration of DNA transposons requires only transposase and no specific host factor, which makes these TEs potentially more successful invaders of diverged genomes (Schaack et al., 2010; Plasterk et al., 1999). The very recent out of Africa habitat expansion of *D. simulans* (Capy and Gibert, 2004) about 100 years ago is therefore consistent with the predominance of bursts of DNA transposon. *D. melanogaster* , on the other hand, colonized already more than 10,000 years ago (Stephan and Li, 2007), providing sufficient time for less invasive retrotransposons to colonize a new host. Furthermore, if *D. melanogaster* experienced a burst of DNA TEs shortly after the colonization, the host defense system (e.g.: the piRNA system (Levin and Moran, 2011)) may have matured to control the initially invading DNA TEs. Under this scenario, the genomic TE signature in *D. simulans* is expected to experience a transition from DNA TE bursts to retrotransposon bursts in the next couple of centuries.

## Materials and Methods

### Fly samples and sequencing

We collected 1,300 isofemale lines of *D. simulans* and 1,250 isofemale lines of *D. melanogaster* from Kanonkop (South Africa) in 2013. The lines were kept in the laboratory for 8 generations. We used a single female from 793 (554) isofemale *D. simulans* (*D. melanogaster* ) lines for pooling. Genomic DNA was extracted from pooled flies using a high salt extraction protocol (Miller et al., 1988) and sheared using a Covaris S2 device (Covaris, Inc. Woburn, MA, USA).

We used three different protocols to prepare paired-end libraries. One library (BGI-91a; supplementary table 1) was prepared following a modified version of the NEBNext Ultra protocol (New England Biolabs, Ipswich, MA). For another library (BGI-92a, BGI-92b, BGI93b; supplementary table 1) we used the NEXTflex PCR-Free DNA Sequencing Kit (Bioo Scientific, Austin, Texas) with modifications. The third library (BGI-93a; supplementary table 1) was prepared based on the NEBNext DNA Sample Prep modules (New England Biolabs, Ipswich, MA) in combination with index adapters from the TruSeq v2 DNA Sample Prep Kit (Illumina, San Diego, CA). All protocols made use of barcoding (supplementary table 1). For each library we selected for a narrow insert size, ranging from 260-340, using agarose gels. A total of five lanes 2x100bp paired-end reads were sequenced on a HiSeq2000 (Illumina, San Diego, CA). In summary we sequenced 364 million paired end fragments for *D. melanogaster* and 288 million paired end fragments for *D. simulans* (supplementary tables 2, 3). This yields an average coverage of 381 in *D. melanogaster* and of 327 in *D. simulans*.

### Annotation of TE insertions

One of the requirements for estimating the abundance of TE insertions with PoPoolation TE (Kofler et al., 2012) is a reliable TE data base. A manually curated high-quality annotation of TE insertions has been generated for *D.melanogaster* (Kaminker et al., 2002; Quesneville et al., 2005) , whereas, to our knowledge, so far no TE annotation of comparable quality exists for *D. simulans*. To avoid any biases that may result from using TE annotations of different qualities we decided to *de novo* annotate TE insertions in both species with an identical pipeline. The reference sequence of *D. melanogaster* (v5.53) was obtained from FlyBase (http://flybase.org). We used the reference sequence published by (Palmieri et al., 2014) for *D. simulans*, as this assembly is of a higher quality than the previous available one (Begun et al., 2007) and of similar quality as a recently published one (Hu et al., 2013). We also obtained a library containing the consensus sequences of *Drosophila* TEs (transposon sequence set.embl; v9.42; (Quesneville et al., 2005)) from FlyBase. To avoid identification of spurious TE insertions we excluded canonical TE sequences not derived from *D. melanogaster* or *D. simulans* (Casey Bergman; personal communication). We mapped the consensus TE sequences against both reference genomes with RepeatMasker open-4.0.3 (Smit et al., 2010) using the RMBlast (v2.2.28) search engine and the settings recommended by (Permal et al., 2012) (-gccalc -s -cutoff 200 -no is -nolow -norna -gff -u), yielding a raw annotation of TE insertions. The consensus sequences of several TE families contain microsatellites which may, as an artefact, be annotated as TE insertions (Permal et al., 2012; Quesneville et al., 2005). To account for this, we identified microsatellites in both reference genomes with SciRoKo 3.4 (Kofler et al., 2007) (required score 12; mismatch penalty 2; seed length 8; seed repeats 3; mismatches at once 3), converted the output into a ’gtf’ file and removed TEs from the raw annotation that overlapped with a microsatellite over more than 30% of the length using bedtools (v2.17.0; intersectBed -a rawannotation.gff -b microsatellites.gff -v -f 0.3) (Quinlan and Hall, 2010). Overlapping TE insertions of the same family were merged and disjoint TE insertions of the same family were linked using an algorithm that, similar to dynamic programming, maximizes the score of the linked TE insertions (*match − score* = 1, *mismatch − penalty* = 0.5). We resolved overlapping TE insertions of different families by prioritizing the longest TE insertion and iteratively truncating the overlapping regions of the next longest insertions. Finally we filtered for TE insertions having a minimum length of 100 bp.

### Estimating the abundance of TE insertions with PoPoolation TE

Estimating the abundance of TE insertions with PoPoolation TE requires paired end sequences from natural populations, a reference sequence, an annotation of TE sequences and a hierarchy of the TE sequences (Kofler et al., 2012). We extracted the hierarchy of TE sequences from the database of consensus TE sequences (v9.42; see above). We extracted the sequences of the annotated TE insertions from the reference genomes into a distinct file and subsequently masked these TE sequences within the reference genome with the character ’N’. We than concatenated the individual fasta records of (i) the consensus sequences of TE insertions, (ii) the TE sequences extracted from the reference genome and (iii) the repeat masked reference genome into a single file, which we call TE-merged-reference. Short read mapping software usually only allows for a few mismatches between read and reference genome which may lead to underestimate the abundance of some TE insertions, especially when the TE sequences are highly diverged (Kofler et al., 2012). Such a high divergence between reads and the reference sequences may also result when the consensus sequences of TE families are derived from a different species. This could lead to underestimate the abundance of TE insertions in *D. simulans* when using consensus sequences that are mostly derived from *D. melanogaster* . Therefore, we improved the sensitivity of our pipeline for *D. simulans* by including TE sequences extracted from the assemblies of Begun et al. (2007), Palmieri et al. (2014) and Hu et al. (2013) (using the same TE annotation pipeline as described above) into the TE-merged-reference of *D. simulans*.

We mapped 364 million PE fragments of *D. melanogaster* and 288 million PE fragments of *D. simulans* (see above) to the respective TE-merged-reference with bwa (v0.7.5a) (Li and Durbin, 2009) using the bwa-sw algorithm (Li and Durbin, 2010) (supplementary tables 2, 3). We used ’samro’ to restore the paired end information (Kofler et al., 2012). We estimated the abundance of TE insertions with PoPoolation TE similarly as described in (Kofler et al., 2012) using the following settings: `identify-te-insertions.pl` –te-hierarchy-level family, –min-count 3, –min-map-qual 15, –narrow-range 100; `crosslink-te-sites.pl` –min-dist 85, –max-dist 300; `estimate-polymorphism.pl` –te-hierarchy-level family, –min-map-qual 15; Subsequently we filtered for TE insertions located on the major chromosome arms (X, 2L, 2R, 3L, 3R, 4) and for TE insertions having a minimum physical coverage of 30 (physical coverage as defined here is the sum of paired end fragments that either confirm the presence or the absence of a TE insertion). An unbiased comparison of the abundance of TE insertions between different species requires similar physical coverages in all species. We therefore iteratively subsampled paired-end fragments and repeated TE identification with PoPoolation TE, until we obtained similar physical coverages in both species (supplementary table 4).

### Estimating nucleotide polymorphism

We estimated genome-wide levels of nucleotide diversity in the two natural populations using Pool-Seq data and PoPoolation (Kofler et al., 2011a). First, we aligned all reads to the respective reference genome (unmodified) with bwa aln (0.7.5a) (Li and Durbin, 2009) and the following parameters: -I -m 100000 -o 1 -n 0.01 -l 200 -e 12 -d 12; Duplicate reads were removed with Picard (v1.95; http://picard.sourceforge.net/). Reads with a mapping quality lower than 20 or reads not mapped as proper pair were removed with samtools (v0.1.19) (Li et al., 2009). We created a pileup file for each population with samtools (v0.1.19) (Li et al., 2009) and the following parameters: -B -Q 0; As alignments spanning indels are frequently unreliable and may lead to spurious SNP calls we removed regions flanking indels (5bp in each direction; minimum count of indel 4) from the pileup with PoPoolation (Kofler et al., 2011a). Subsequently we subsampled the pileup to a uniform coverage of 175 with PoPoolation (Kofler et al., 2007) and the following parameters: –maxcoverage 1400 –min-qual 20 –method withoutreplace; Finally we calculated *π* for windows of 100kb with PoPoolation and the following paramters: –min-count 4 –min-coverage 165 –maxcoverage 175 –min-covered-fraction 0.6 –min-qual 20 –no-discard-deletions –pool-size 1300; For estimating *F*_*ST*_ between *D. simulans* populations we used PoPoolation2 (Kofler et al., 2011b) and the following parameters –min-qual 20 –window-size 1 –step-size 1 –pool-size 500–suppress-noninformative –min-covered-fraction 1 –max-coverage 600,100,100 –min-count 4–min-coverage 10; Only autosomes were used for calculating the *F*_*ST*_ .

### Expression level of transposable element families in D.simulans

To measure the expression level of different TE families in *D. simulans* we obtained previously published RNA-seq reads (Palmieri et al., 2014), derived from a mix of several developmental stages of *D. simulans* strain M252. The reads were trimmed with PoPoolation v1.2.2 (trim-fastq.pl) (Kofler et al., 2011a) using the following parameters: –fastq-type illumina, –quality-threshold 20, –min-length 40; We mapped the RNA-seq reads to a database consisting of the repeat masked reference genome of *D. simulans* (Palmieri et al., 2014) and the library of TE sequences derived from all three assemblies of *D. simulans* (see above). Reads were mapped with bwa (v0.7.5a) (Li and Durbin, 2009) using the bwa-sw algorithm (Li and Durbin, 2010). Unambiguously mapped reads (mapping quality ≥ 15) were filtered with samtools (v0.1.19) (Li et al., 2009). Subsequently we counted the number of reads mapping to each TE family and normalized counts by the length of the consensus sequence (transposon sequence set.embl; v9.42; see above).

### Orthologous regions between D. melanogaster and D. simulans

The assemblies of *D. melanogaster* and *D. simulans* are of different quality, for example varying in the amount of assembled heterochromatin. An unbiased analysis of TE abundance should therefore be restricted to genomic regions being present in the assemblies of both species. We identified these regions by aligning the genomes of *D. melanogaster* (v5.53) and *D. simulans* (Palmieri et al., 2014) with MUMmer (v3.23; nucmer) (Kurtz et al., 2004). To avoid spurious alignments we masked all sequences derived from TEs in both reference genomes (see above) prior to the alignment. Coordinates were extracted with the ’show-coords’ tool (Kurtz et al., 2004) and only alignments of the major chromosome arms (X, 2L, 2R, 3L, 3R, 4) were considered. Due to the masking of TE sequences these raw alignments contain a plenitude of gaps where the TE insertions actually causing the gaps, are not found in genomic regions that are present in the alignment. To mitigate this we linked these gaps by merging alignments not separated by more than 20.000bp in both species. This threshold of 20.000bp has been arbitrary chosen because only six of the masked regions in the repeat-masked genome of *D. melanogaster* have a size larger than 20.000bp.

### Modeling TE abundance in populations under an equilibrium model

We performed forward simulations for estimating the variance of TE abundance in natural populations expected under an equilibrium model. The simulations aimed to capture conditions found in *D. melanogaster* and accordingly we (i) simulated diploid organisms, (ii) used a genome with a similar size and number of chromosomes as *D. melanogaster* and (iii) used the recombination rate of *D. melanogaster* . We obtained the recombination rate from the *D. melanogaster* recombination rate calculator v2.2 (Fiston-Lavier et al., 2010) for windows of 1000kb. We excluded the X-chromosome and low recombining regions including the entire chromosome 4 from the analysis. In summary we performed our simulations with *T* = 68,700,000 TE insertions sites (distributed over the following genomic regions 2L:300000-16600000, 2R:3900000-20700000, 3L:900000-17400000, 3R:6600000-25700000) where every insertion site may either be empty or occupied. In our model, every TE insertion has a constant probability of transposing to a novel site *v* and excision events (*u* = 0) were not considered. Novel TEs were randomly inserted in any of the *T* insertion sites at any of the two haploid genomes. If an insertion site was already occupied the transposition event was ignored. For any individual *i* in a population of size *N* the fitness *w*_*i*_ can be calculated as 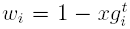, where *g*_*i*_ is the number of TE insertions, *x* is the selective disadvantage of each insertion and *t* represents the interactions between the insertions (Charlesworth and Charlesworth, 1983). This is a model where all TE insertions exert a semi-dominant effect (Charlesworth and Charlesworth, 1983).

Per default we used *x* = 0.0004 and *t* = 1.3 in our simulations. We furthermore used fecundity selection, where any individual has a probability of mating *p*_*i*_ that linearly scales with fitness 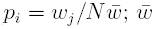 is the average fitness; after Le Rouzic et al. (2007)).

We simulated evolving populations with non-overlapping generations, proceeding at every generation in the following order: First *N* random pairs were picked according to the mating probability *p*_*i*_, where selfing was excluded. Second, each parent contributed a single gamete to the offspring wherein crossing over events were introduced according to the specified recombination rate (see above). Third, fitness of the offspring *w*_*i*_ was calculated from the abundance of TE insertions in the resulting genome of the offspring. And fourth, transposition events were introduced according to the transposition rate *v*. Note that the novel TE insertions will only contribute to fitness in the next generation. This could for example be interpreted as TE activity in the germline which will mostly also only effect the next generation (i.e.: the offspring). In all simulations, we performed forward simulations for 10,000 generations. We noted that if a stable equilibrium could be reached (e.g.: no increase in the number of fixed insertions), it took less than 5,000 generations. Conservatively we used 10,000 generations in our simulations. To match the analysis of natural populations we also sampled 145 haploid genomes after the 10,000 generations and required a minimum count of 3 to identify a transposable element (see above).

#### Constant population size

In order to estimate the expected variance in TE copy number under an equilibrium model and an constant population size, we performed forward simulations for populations of *N* = 10.000 diploid individuals. We performed 10,427 individual forward simulations with transpositions rates randomly sampled from a uniform distribution between *v* = 0.0 *−* 0.003. These simulations required approximately 10,000 CPU hours. Different TE families may have markedly different transposition rates (Charlesworth and Langley, 1989) which will result in different equilibrium copy numbers. We therefore identified for every TE family (*j*) the most likely transposition rate *v* that maximizes the probability of observing both the TE copy number of *D. melanogaster* 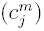 and of *D. simulans* 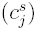. To do this, we grouped the simulation results based on the transposition rate *v* into *i* overlapping windows (*W*_*i*_ ∈ *W*) with a window size of 10^−4^ and a step size of 10^−5^ and fitted, for every window, a normal distribution to the data 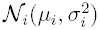 with mean *μ_i_* and standard deviation 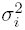). The probability that a given number of TE insertions (*c*) can be explained by the transposition rate of window *W*_*i*_ is than given by *P* (*c|W_i_*) = 1 *− P* (*μ_i_ − |μ_i_ − c| < x < μ_i_* + *|μ_i_ − c|*) which can be easily computed from \*\*\*\*\**N*_*i*_.

Next we identified for every TE family (*j*) the window 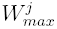 that maximizes the probability of observing 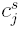 and 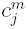 as 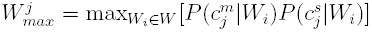. The corresponding transposition rate of this window will also be the maximum likelihood estimate of *v*. Finally the probability of observing both 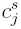 and 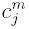 with a constant transposition rate as found in window 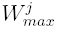 can be computed as 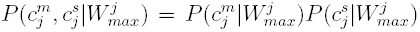. We tested every TE family for significance using Bonferroni correction to account for multiple testings.

#### Varying population size

In order to include demography into our model of TE dynamics we estimated differences in effective population sizes by comparing the level of nucleotide polymorphism in *D. melanogaster* and *D. simulans*. We found that *D. simulans* has a 1.519 higher effective population size than *D. melanogaster* . Accordingly, we performed forward simulations with two different population sizes where the larger population (*N* = 10.000) represents *D. simulans* and the smaller population (*N* = 6,583; *≈* 10000/1.519) represents *D. melanogaster* . Differences in TE insertions between these two species were assessed as described above. The only difference was that, for every window (*i*) we fitted two separate normal distriubtions to the data, one for *D. melanogaster* 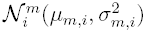 and one for *D. simulans* 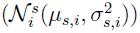. The probability that a given number of TE insertions in *D. melanogaster* (*c*^*m*^) can be explained by the transposition rate of the given window (*W*_*i*_) can be calculated as 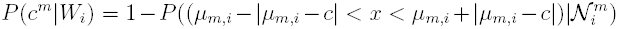, and accordingly, the probability that the number of TE insertions in *D. simulans* can be explained by the transposition rate in the same window as 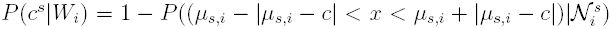. Finally, the maximum likelihood window and the probability of observing both TE counts with the window-specific transposition rate were computed as described above. Again, we used Bonfferoni correction to account for multiple testing.

### Data availability

The short reads (European Nucleotide Archive; http://www.ebi.ac.uk/ena; PRJEB6673) and the TE annotations (https://code.google.com/p/popoolationte/wiki/pdms) are publicly available. All scripts and the entire protocol used for this work ar also available (https://code.google.com/p/popoolationte/wiki/pdms)

## Acknowledgments

We thank all members of the Institute of Population Genetics for feedback and support. We especially thank Casey Bergman for valuable advice. We are grateful to Hester van Schalkwyk for fly collection and to Núria Ortiz Mompel, Ariana Macon, Sofia Lehrner, Stephanie Lilja, Siegrid Widhalm and Renate Weiß for expert fly work. This work was supported by the ERC grant Archadapt and Austrian Science Funds (FWF) grant P22725.

## 1 Supplementary Tables

**Table 1:**
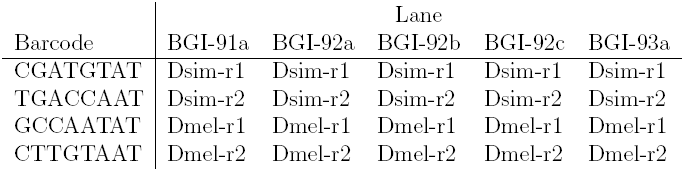
Sample IDs for the barcodes used in the sequenced Illumina paired-end lanes; Dsim: *D. simulans*; Dmel: *D. melanogaster* ; r1: replicate 1; r2: replicate 2

| Barcode | Lane |  |  |  |  |
| --- | --- | --- | --- | --- | --- |
|  | BGI-91a | BGI-92a | BGI-92b | BGI-92c | BGI-93a |
| CGATGTAT | Dsim-r1 | Dsim-r1 | Dsim-r1 | Dsim-r1 | Dsim-r1 |
| TGACCAAT | Dsim-r2 | Dsim-r2 | Dsim-r2 | Dsim-r2 | Dsim-r2 |
| GCCAATAT | Dmel-r1 | Dmel-r1 | Dmel-r1 | Dmel-r1 | Dmel-r1 |
| CTTGTAAT | Dmel-r2 | Dmel-r2 | Dmel-r2 | Dmel-r2 | Dmel-r2 |

**Table 2:**
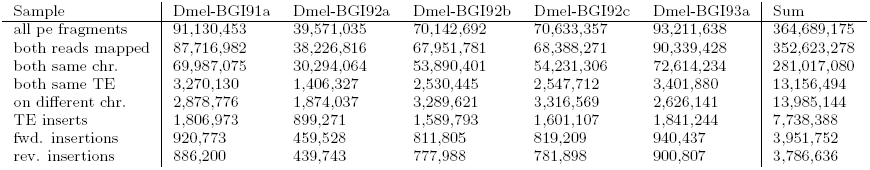
Mapping statistics for *D. melanogaster* ; All numbers are counts of paired end (pe) fragments; both same chr.: both reads mapped to the same reference chromosome; both same TE: both reads mapped to the same TE; on different chr.: reads mapped to different reference chromosomes; TE inserts: one read maps to a TE and the other to a reference chromosome; fwd. insertions: forward insertions; rev. insertions: reverse insertions for details see (Kofler et al., 2012)

**Table 3:** Mapping statistics for *D. simulans*; All numbers are counts of paired end (pe) fragments; both same chr.: both reads mapped to the same reference chromosome; both same TE: both reads mapped to the same TE; on different chr.: reads mapped to different reference chromosomes; TE inserts: one read maps to a TE and the other to a reference chromosome; fwd. insertions: forward insertions; rev. insertions: reverse insertions for details see (Kofler et al., 2012)

| Sample | Dsim-BGI91a | Dsim-BGI92a | Dsim-BGI92b | Dsim-BGI92c | Dsim-BGI93a | Sum |
| --- | --- | --- | --- | --- | --- | --- |
| all pe fragments | 69,083,283 | 27,637,652 | 53,590,534 | 53,682,148 | 84,513,925 | 288,507,542 |
| both reads mapped | 57,948,905 | 24,279,922 | 47,108,790 | 47,180,635 | 71,705,665 | 248,223,917 |
| both same chr. | 48,352,523 | 19,849,495 | 38,477,056 | 38,538,543 | 59,940,332 | 205,157,949 |
| both same TE | 2,080,323 | 642,041 | 1,253,782 | 1,254,677 | 2,559,287 | 7,790,110 |
| on different chr. | 918,263 | 1,108,546 | 2,151,003 | 2,155,971 | 1,205,970 | 7,539,753 |
| TE inserts | 1,501,665 | 688,985 | 1,347,272 | 1,346,956 | 1,793,507 | 6,678,385 |
| fwd. insertions | 755,392 | 347,166 | 678,845 | 679,123 | 900,686 | 3,361,212 |
| rev. insertions | 746,273 | 341,819 | 668,427 | 667,833 | 892,821 | 3,317,173 |

**Table 4:** Mapping statistics after subsampling of paired end fragments; All numbers are counts of paired end (pe) fragments (except for res. phys. cov.); both same chr.: both reads mapped to the same reference chromosome; both same TE: both reads mapped to the same TE; on different chr.: reads mapped to different reference chromosomes; TE inserts: one read maps to a TE and the other to a reference chromosome; fwd. insertions: forward insertions; rev. insertions: reverse insertions; res. phys. cov.: resulting average physical coverage of a TE insertion; for details see (Kofler et al., 2012)

|  | *D.melanogaster* | *D. simulans* |
| --- | --- | --- |
| all pe fragments | 306,813,382 | 288,507,542 |
| both reads mapped | 296,928,194 | 248,223,917 |
| both same chr. | 236,625,431 | 205,157,949 |
| both same TE | 11,081,095 | 7,790,110 |
| on different chr. | 12,133,864 | 7,539,753 |
| TE inserts | 6,587,515 | 6,678,385 |
| fwd. insertions | 3,365,526 | 3,361,212 |
| rev. insertions | 3,221,989 | 3,317,173 |
| res. phys. cov. | 144.986 | 145.821 |

**Table 5:**
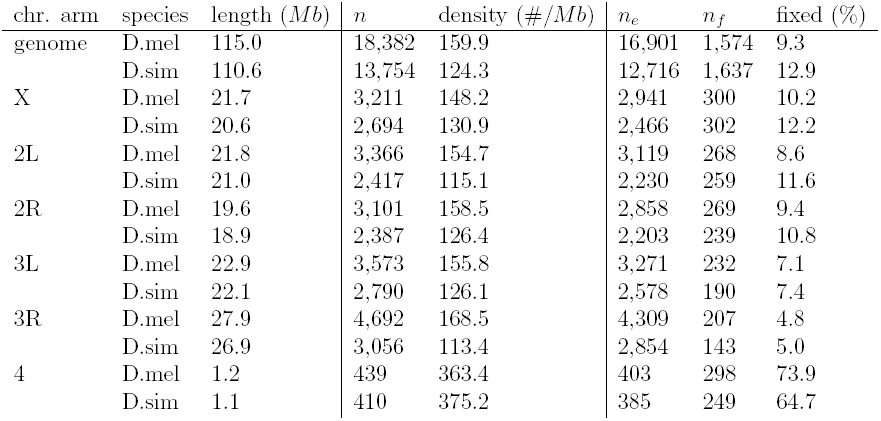
The abundance of TE insertion in *D. melanogaster* (D.mel) and *D. simulans* (D.sim). Only TE insertions in genomic regions being present in the assemblies of both species are considered. *n* number of TE insertions; *n*_*e*_ number of TE insertions for which population frequencies could be estimated (not overlapping, minimum physical coverage of 30); *n*_*f*_ number of fixed insertions; chr. arm: chromosome arm

| chr. arm | species | length (Mb) | $n$ | density (#/Mb) | $n_e$ | $n_f$ | fixed (%) |
| --- | --- | --- | --- | --- | --- | --- | --- |
| genome | D.mel | 115.0 | 18,382 | 159.9 | 16,901 | 1,574 | 9.3 |
|  | D.sim | 110.6 | 13,754 | 124.3 | 12,716 | 1,637 | 12.9 |
| X | D.mel | 21.7 | 3,211 | 148.2 | 2,941 | 300 | 10.2 |
|  | D.sim | 20.6 | 2,694 | 130.9 | 2,466 | 302 | 12.2 |
| 2L | D.mel | 21.8 | 3,366 | 154.7 | 3,119 | 268 | 8.6 |
|  | D.sim | 21.0 | 2,417 | 115.1 | 2,230 | 259 | 11.6 |
| 2R | D.mel | 19.6 | 3,101 | 158.5 | 2,858 | 269 | 9.4 |
|  | D.sim | 18.9 | 2,387 | 126.4 | 2,203 | 239 | 10.8 |
| 3L | D.mel | 22.9 | 3,573 | 155.8 | 3,271 | 232 | 7.1 |
|  | D.sim | 22.1 | 2,790 | 126.1 | 2,578 | 190 | 7.4 |
| 3R | D.mel | 27.9 | 4,692 | 168.5 | 4,309 | 207 | 4.8 |
|  | D.sim | 26.9 | 3,056 | 113.4 | 2,854 | 143 | 5.0 |
| 4 | D.mel | 1.2 | 439 | 363.4 | 403 | 298 | 73.9 |
|  | D.sim | 1.1 | 410 | 375.2 | 385 | 249 | 64.7 |

## 2 Supplementary figures

**Figure 1:**
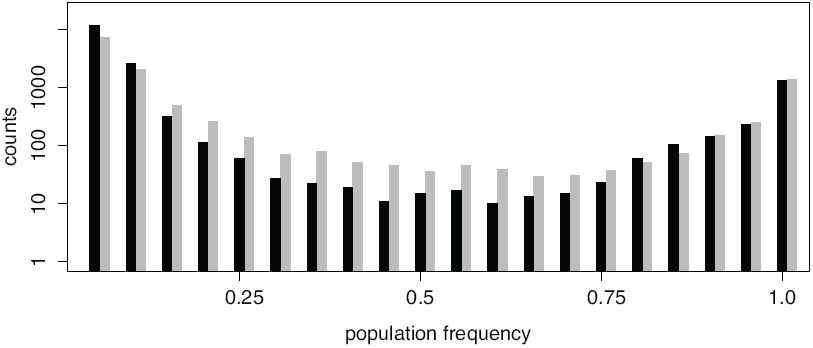
Frequency distributions of TE insertions in *D. melanogaster* (black) and *D. simulans* (grey); Only
TE insertions for which the population frequencies could be estimated are shown (not overlapping, minimum physical coverage of 30); *D. melanogaster*: 16; 901 insertions; *D. simulans*: 12; 716 insertions

**Figure 2:**
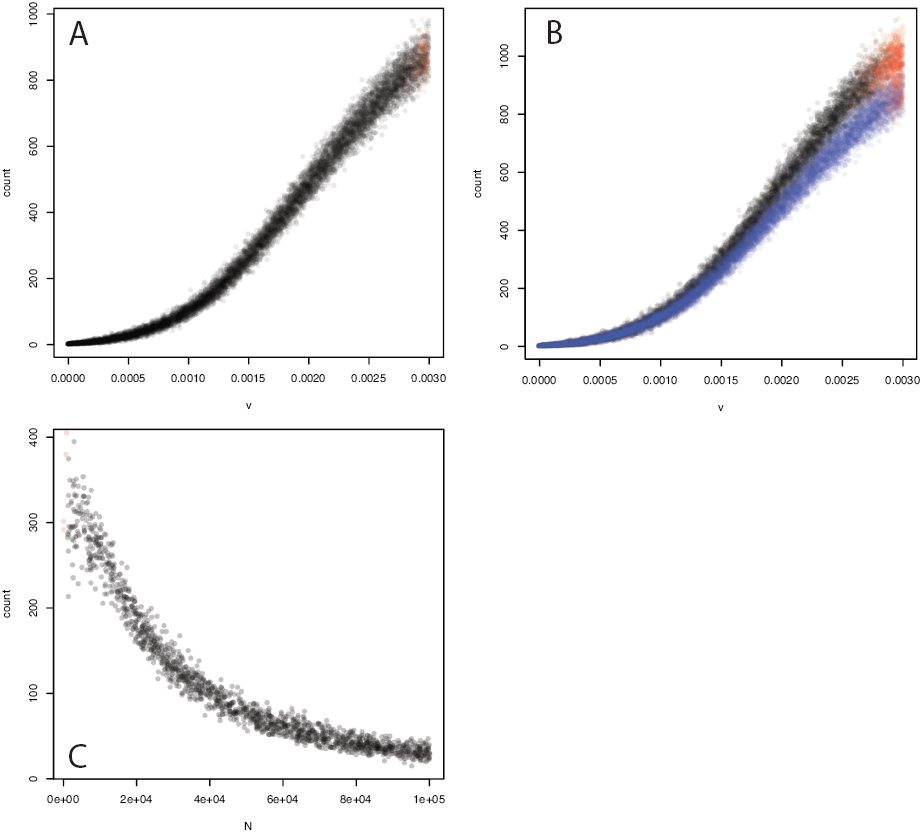
Simulations of TE abundance in populations under an equilibrium model. A) Equilibrium copy numbers of TE insertions for a population of size *N* = 10,000 (black dots). In total 10,427 forward simulations have been performed for randomly drawn transposition rates (*v*) ranging from 0.0 to 0.003. B) Equilibrium copy numbers of TE insertions for populations sizes of *N* = 6,583 (black dots) and *N* = 10,000 (blue dots). C) Influence of the population size. A fixed transposition rate of *v* = 0.0015 was used and 1,664 forward simulations have been performed for randomly drawn population sizes (*N* ) ranging from 100 to 100,000. Equilibrium copy numbers decrease with increasing population size. In all experiments (A,B,C) we simulated TE dynamics for 10,000 generations. Red dots indicate, irrespective of *N* , populations with more than 5 fixed TE insertions. These populations will, in the course of time, accumulate increasing numbers of fixed TE insertions and are therefore gradually loosing the ability to maintain TE copy numbers at a stable equilibrium. For more details about the simulations see material and methods of the main manuscript

**Figure 3:**
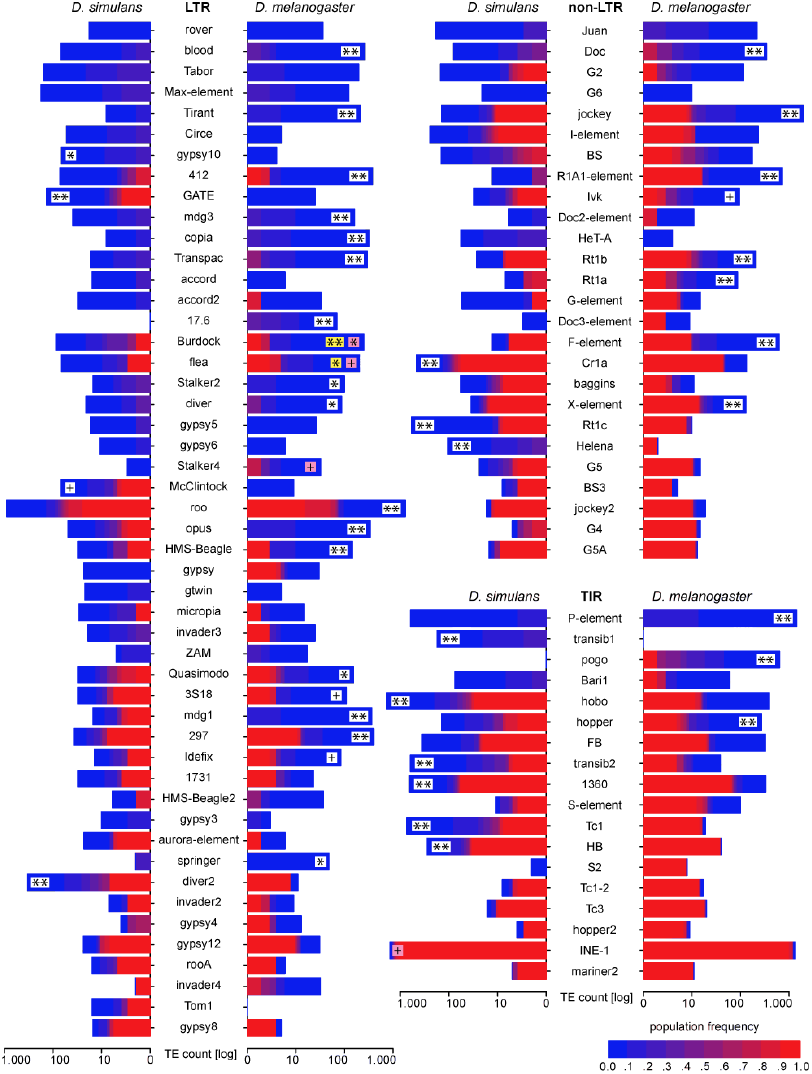
Abundance of different TE families in natural *D. melanogaster* and *D. simulans* populations; Significant differences in TE copy numbers are indicated for the species with a higher number of insertions, assuming equal population sizes in both species (yellow), or a *N*_*e*_ ratio of 1.519 (pink). Those cases for which both models agree are indicated in white. p-value after Bonferroni correction: ** < 0.001; * < 0.01; + < 0.05; Only TE families having in total more than 10 insertions are shown

## 3 Supplementary Results

### 3.1 Sensitivity and specificity of the TE annotation pipeline

To test the performance of our pipeline for the *de novo* annotation of TE insertions we used the reference annotation of *D. melanogaster* (FlyBase v5.53) as ’gold standard’ and asked whether our pipeline reproduces this reference annotation. We excluded peri-centromeric regions that have, so far, not been annotated for TE insertions (2R:>22,420,241bp, 2L:<387,345bp, 3L:>23,825,333bp; Casey Bergman personal communication). We found that our *de novo* annotation pipeline has a high sensitivity and specificity both at the nucleotide level and the level of individual TE insertions (table 6).

### 3.2 Quality control for species pools

We used a very large number of individuals (> 500) from both species to establish isofemale lines and subsequently the pools used for this study. Since *D. melanogaster* and *D. simulans* are phenotypically similar, two different people checked each isofemale line. Since, it is possible that an error occurred in the species identification, we decided to additional check the sequenced pools. We compiled a set of 9,491 SNPs on chromosome 4 that are fixed for different alleles in the two species (R. Tobler, pers. communication). Using these SNPs we found that 0.042% of the base calls in the *D. melanogaster* library are identical to the allele fixed in *D. simulans*, which is close to the fraction of sequencing errors in this library (0.035%). Similarly for the *D. simulans* library we found that 0.014% of the base calls are identical to the allele fixed in *D. melanogaster* , which is again similar to the level of sequencing errors in this library (0.018%). The level of sequencing errors was estimated as the fraction of base calls at these 9,491 SNPs that are neither identical to the allele fixed in *D. melanogaster* nor to the allele fixed in *D. simulans*. We therefore conclude that each of the pools of individuals was derived from a single species only.

**Table 6:** Performance of the *de novo* annotation pipeline for TE insertions relative to the reference annotation of *D. melanogaster* (*D.mel.* v5.53). The performance was measured at the level of individual insertions (number of insertions having at least one nucleotide overlap; ins.) and at the nucleotide level (nuc). We estimated the number of true positives (TP) and of false positives (FP). Numbers in brackets are percentages relative to the reference annotation.

|  | *D.mel.* v5.53 | TP | FP |
| --- | --- | --- | --- |
| ins. | 5,432 | 4,516 (83.1%) | 534 (9.8%) |
| nuc. | 6,556,993 | 6,266,442 (95.6%) | 343,628 (5.2%) |

### 3.3 Reproducibility of results of Kofler et al. (2012)

We also tested whether estimates of TE abundance generated with our *de novo* TE annotation match previously published results, that were generated with the reference annotation (Kofler et al., 2012). Kofler et al. (2012) estimated the TE abundance in a natural population of *D. melanogaster* from northern Portugal (Povoa de Varzim) using PoPoolation TE and the reference annotation. We compared the estimates of TE abundance obtained in this work with the results of Kofler et al. (2012) and found a good agreement for the population frequency (fig. 4A; Spearman’s rank correlation, *r*_*S*_ = 0.82, *p* < 2.2*e* − 16) as well as for the number of insertions (fig. 4B; Spearman’s rank correlation, *r*_*S*_ = 0.81, *p* < 2.2*e* − 16). We note that some deviation in the TE abundance between these two samples are expected (Vieira et al., 1999). This good agreement between estimates of TE abundance despite different annotations, different geographic origins of the populations, different read length (here 100 vs 74), different library preparation methods suggests that our approach yields highly reliable estimates of TE abundance. In agreement with this, the reliability of PoPoolation TE was recently also confirmed by a simulation study (Zhuang et al., 2014). However, some TE families show marked differences in the numbers of TE insertions between Portugal and South Africa (fig. 4). While the lack of P-element insertions in the population from Portugal can simply be explained by the fact that P-elements were not considered in the study of Kofler et al. (2012), the higher copy numbers of R1A1-elements (South Africa 746; Portugal 11) in South Africa and of gypsy2 (South Africa 14; Portugal 50) elements in Portugal, may be due to different activities of these TE families in the two populations.

### 3.4 Reproducibility of results of Vieira et al. (1999)

Vieira et al. (1999) estimated the abundance of 36 TE families in multiple populations of *D. melanogaster* and *D. simulans* using *in situ* hybridization. In order to enable comparing our data with the results of Vieira et al. (1999), who provided the abundance of TE families as average counts per individual genome, we simply weighted every TE insertion by it’s population frequency (population frequency can be interpreted as the probability of observing a given insertion in a random genome). We did not include TE families for which Vieira et al. (1999) and our study, did not find a single insertion (*osvaldo*, *gandalf*, *telemac*, *bilbo*), as inclusion of such families may lead to artificially inflated correlations. We also did not include P-element and mariner insertions, as abundance of these two families were not directly estimated by Vieira et al. (1999). Overall we found a striking correlation between TE abundance estimated in this study and the study of Vieira et al. (1999), both for *D. melanogaster* (Spearman’s rank correlation; *r*_*S*_ = 0.85, *p* = 3.6*e* − 9) and *D. simulans* (Spearman’s rank correlation; *r*_*S*_ = 0.62, *p* = 0.0002). We note that absolute insertion numbers cannot be compared directly, as Vieira et al. (1999) excluded pericentromeric regions and provided the TE abundance per diploid genome, whereas we only analyzed regions being present in the assemblies of both species and provided the TE abundance per haploid genome.

**Figure 4:**
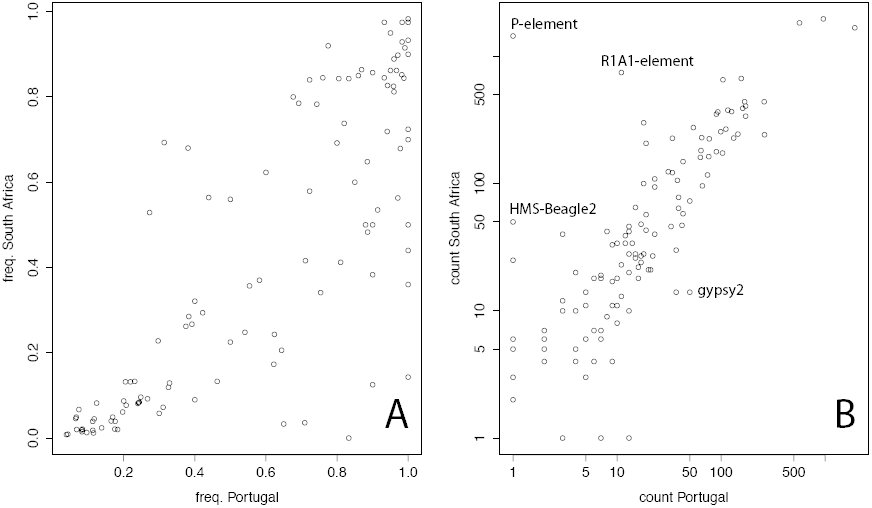
Comparison of the abundance of TE families in two natural populations of *D. melanogaster* : a population from northern Portugal (from Kofler et al. (2012)) and a population from Southern Africa (this work). The average population frequency (A) and the number of insertions (B) are shown. freq.: frequency

**Table 7:** Number of TE insertions per genome for natural populations of *D. melanogaster* (*D.mel.*) and *D. simulans* (*D.sim.*) as estimated by Vieira et al. (1999) and by this study.

| family | Vieira et al. (1999) |  | this study |  |
| --- | --- | --- | --- | --- |
|  | *D.mel.* | *D.sim.* | *D.mel.* | *D.sim.* |
| 1731 | 1.55 | 1.00 | 5.4 | 5.8 |
| 17.6 | 2.50 | 0.00 | 7.3 | 0.0 |
| 297 | 23.40 | 1.00 | 40.5 | 18.0 |
| 412 | 28.45 | 13.88 | 16.0 | 6.8 |
| BEL/3S18 | 5.25 | 0.58 | 7.7 | 9.2 |
| blood | 17.45 | 2.5 | 10.35 | 4.3 |
| burdock | 10.35 | 5.27 | 13.0 | 10.3 |
| copia | 24.05 | 3.88 | 13.4 | 0.7 |
| coral/transpac | 15.85 | 1.88 | 13.8 | 1.8 |
| flea | 16.60 | 3.42 | 16.0 | 9.3 |
| gypsy | 1.70 | 1.54 | 7.2 | 1.9 |
| HMS beagle | 9.50 | 2.77 | 9.3 | 6.8 |
| idefix | 5.70 | 1.00 | 10.8 | 6.3 |
| mdg1 | 20.75 | 0.19 | 16.4 | 7.3 |
| mdg3 | 14.10 | 3.35 | 7.8 | 4.4 |
| opus | 20.90 | 4.81 | 18.1 | 8.8 |
| prygun/rover | 11.35 | 0.81 | 4.2 | 0.9 |
| roo | 67.60 | 38.46 | 122.6 | 100.4 |
| springer | 2.35 | 0.00 | 2.7 | 1.3 |
| stalker | 6.50 | 0.38 | 1.7 | 1.0 |
| tirant | 11.45 | 1.62 | 7.8 | 0.9 |
| ZAM | 0.35 | 0.23 | 1.5 | 1.7 |
| Doc | 26.20 | 13.81 | 20.3 | 7.7 |
| F-element | 31.40 | 1.77 | 36.3 | 8.3 |
| helena | 0.25 | 10.23 | 2.8 | 12.9 |
| I | 25.15 | 12.58 | 10.6 | 33.7 |
| jockey | 31.60 | 3.27 | 93.7 | 21.4 |
| bari-1 | 4.37 | 4.88 | 8.2 | 4.5 |
| hobo | 49.90 | 66.23 | 29.9 | 136.4 |
| pogo | 13.25 | 0.00 | 39.5 | 0.0 |

**Table 8:** Nucleotide polymorphism in the natural *D. melanogaster* (*D.mel.*) and *D. simulans* (*D.sim.*) populations. From genomic regions being present in both assemblies (both assemblies) we successively removed low recombining regions (<1 cM/Mb; excl. low. rec) and regions overlapping with exons (excl. coding).

|  | both assembl. |  | excl. low. rec. |  | excl. coding |  |
| --- | --- | --- | --- | --- | --- | --- |
|  | *D.mel.* | *D.sim.* | *D.mel.* | *D.sim.* | *D.mel.* | *D.sim.* |
| whole genome | 0.0061 | 0.0104 | 0.0068 | 0.0108 | 0.0070 | 0.0107 |
| X | 0.0052 | 0.0077 | 0.0053 | 0.0082 | 0.0056 | 0.0086 |
| 2L | 0.0072 | 0.0115 | 0.0080 | 0.0121 | 0.0081 | 0.0119 |
| 2R | 0.0063 | 0.0108 | 0.0070 | 0.0109 | 0.0070 | 0.0107 |
| 3L | 0.0067 | 0.0116 | 0.0078 | 0.0118 | 0.0080 | 0.0117 |
| 3R | 0.0055 | 0.0108 | 0.0064 | 0.0112 | 0.0067 | 0.0108 |
| 4 | 0.0012 | 0.0014 | - | - | - | - |
| autosomes | 0.0063 | 0.0110 | 0.0072 | 0.0115 | 0.0074 | 0.0113 |

### 3.5 Nucleotide polymorphism

We estimated the nucleotide polymorphism in the natural *D. melanogaster* and *D. simulans* populations with a sliding window approach, using non-overlapping windows of 1 kb (see material and methods in main manuscript). Subsequently we (i) filtered for regions being present in both assemblies, (ii) excluded regions with low recombination rates (< 1 *cM/M b*) in *D. melanogaster* and (iii) excluded windows overlapping with an exon (table 8). Based on the level of polymorphism in the autosomes we estimate that the *N*_*e*_ of *D. simulans* is approximately 1.519 (= 0.011285/0.007429) times higher than the *N*_*e*_ of *D. melanogaster* .

### 3.6 Age of fixed TE insertions

Reasoning that fixed TE insertions are mostly old, we asked whether fixed insertions are enriched for insertions shared between *D. melanogaster* and *D. simulans*. Shared insertions should mostly predate the split of these two species, which occurred approximately 2-3 million years ago (Hey and Kliman, 1993; Lachaise et al., 1988). To do this, we first generated a set of TE insertions that may potentially be shared between these two species by reciprocally aligning 1000*bp* regions flanking the TE insertion, both at the 5’ and the 3’ end, to the reference genomes of *D. melanogaster* and *D. simulans* using bwa-sw (v0.7.5a) (Li and Durbin, 2010). We only retained TE insertions (i) where the flanking regions could be unambiguously mapped (mapping quality ≥ 15) and (ii) where the flanking regions could be mapped back to the original positions. This procedure yielded a set of 15,079 TE insertions that are potentially shared between the two species, i.e.: insertions in non-repetitive regions and insertions in regions that are present in the assemblies of both species (table 9). Actually shared insertions where subsequently identified by scanning for TE insertions of a given family having, in both species, insertion positions within the boundaries of these flanking regions. Note that this procedure allows for some degree of uncertainty in the exact insertion position of the TEs (as for example advisable when using PoPoolation TE). We generated an additional data set excluding the P-element, which has a strong insertion bias (Spradling et al., 2011) that may potentially bias our results. To ensure that any significant enrichment is not solely based on INE-1, a very old TE family that has not been active for >3 million years (Kapitonov and Jurka, 2003; Sackton et al., 2009; Singh and Petrov, 2004), we also generated a data set excluding INE-1 insertions (table 9).

**Table 9:** Numbers of fixed (fix) or segregating (nfix) TE insertions being present either in both species (sh) or just in *D. melanogaster* (nsh). Data are shown for all insertions that could potentially be shared between both species (all), and subsets of these data, excluding either only the P-element (excl. P) or the P-element and INE-1 (excl. P,I).

|  | all | excl. P | excl. P,I |
| --- | --- | --- | --- |
| fix-sh | 577 | 577 | 57 |
| fix-nsh | 188 | 188 | 100 |
| nfix-sh | 868 | 440 | 315 |
| nfix-nsh | 13,446 | 12,491 | 12,378 |
| sum | 15,079 | 13,696 | 12,850 |

We found that, for all three data sets (table 9), fixed TE insertions are indeed enriched for shared insertions (Fisher’s exact test; *p* < 2.2*e* − 16).

### 3.7 TE families evolving according to the equilibrium or non-equilibrium model

The equilibrium model requires constant activity of a given TE family for some extended period of time. Every family deviating from this pattern, by for example showing only past or only recent activity, is therefore evolving according to a non-equilibrium model. In the manuscript we suggested the following approximation: TE insertions having low population frequency are mostly young whereas fixed TE insertions are mostly old. Using this simplification we can roughly estimate whether the TE families shown in supplementary fig. 3 evolve according to an equilibrium or a non-equilibrium model.

#### 3.7.1 only present in one species; non-equilibrium

To this category we assign families that are only present in one species. (supplementary fig. 3).

- LTR (2): 17.6, Tom1
- TIR (3): transib1, pogo, S2

#### 3.7.2 only recent activity in at least one species; non-equilibrium

To this group we assign families that are only active since very recently in at least one species. No past activity, either in one or in both species, could be detected. With respect to supplementary fig. 3 this translates to families having just blue insertions (young: low population frequency) in at least one species.

- LTR (27): rover, blood, Tabor, Max-element, Tirant, Circe, gypsy10, GATE, mdg3, copia, Transpac, accord, accord2, Stalker2, diver, gypsy5, gypsy6, Stalker4, McClintock, opus, gypsy, gtwin, invader3, ZAM, mdg1, gypsy3, springer
- non-LTR (7): Juan, Doc, G6, Helena, Doc2-element, HeT-A, Doc3-element
- TIR (2): P-element, Bari1

#### 3.7.3 3.7.3 inactive in at least one species; non-equilibrium

To this group we assign families that are inactive in at least one species. Allowing for some small margin of methodological error (PoPoolation TE may, due to false absence reads, occasionally underestimate the population frequency of TE insertions (Kofler et al., 2012)), this translates to families having mostly red (old: fixed) insertions in at least one species (supplementary fig. 3).

- LTR (3): invader4, gypsy8, rooA
- non-LTR (4): BS3, jockey2, G4, G5A
- TIR (5): Tc1-2, Tc3, hopper2, INE-1, mariner2

#### 3.7.4 3.7.4 different activity in the two species; non-equilibrium

To this group we assign families that were active for some time (having old and young insertions) and that furthermore have copy numbers in the two species which significantly deviate, mostly because of young insertions, from expectations under an equilibrium model. With respect to supplementary fig. 3 this translates to families having red and blue insertions in the two species, but with significantly different copy numbers.

- LTR (1): 412, Burdock, flea, roo, HMS-Beagle, Quasimodo, 3S18, 297, Idefix, diver2
- non-LTR (9): jockey, R1A1-element, Ivk, Rt1b, Rt1a, F-element, Cr1a, X-element, Rt1c
- TIR (6): hobo, hopper, transib2, 1360, Tc1, HB

#### 3.7.5 3.7.5 similar activity in both species; equilibrium

To this group we assign families that were active for some time (having old and young insertions) and that furthermore have copy numbers in the two species **not** significantly deviating from expectations under an equilibrium model. This translates to families having similar amounts of blue and red insertions in both species (supplementary fig. 3).

- LTR (7): micropia, 1731, HMS-Beagle2, aurora-element, invader2, gypsy4, gypsy12
- non-LTR (6): G2, I-element, BS, G-element, baggins, G5
- TIR (2): FB, S-element

### 3.8 TE families showing signatures of recent bursts

Using the approximation suggested in the manuscript TE insertions having low population frequency are mostly young whereas fixed insertions are mostly old we can roughly distinguish between two different types of bursts of TE activity.

1.) A novel TE family, that was for example horizontally transferred (Bartolomé et al., 2009; Loreto et al., 2008; Sánchez-Gracia et al., 2005), may immediately have a marked activity. We regard the presence of several young insertions combined with the absence of old insertions as signature of such a burst of a novel family (supplementary fig. 3).

2.) Alternatively an extant TE family that had a constant activity for an extended period of time, may show a sudden marked increase in activity. We regard the presence of both old and young insertions combined with significantly different copy numbers in the two species as signature of such a burst of a previously dormant family (supplementary fig. 3).

#### 3.8.1 3.8.1 burst of novel family in D. simulans

- LTR (23): rover, blood, Tabor, Max-element, Tirant, Circe, gypsy10, mdg3, copia, Transpac, accord, accord2, Stalker2, diver, gypsy5, gypsy6, Stalker4, gypsy, gtwind, invader3, ZAM, gypsy3, springer
- non-LTR (8): Juan, Doc, G6, R1A1-element, Doc2-element, HeT-A, Doc3-element, Helena
- TIR (4): P-element, transib1, Bari1, S2

#### 3.8.2 burst of novel family in D. melanogaster

- LTR (24): rover, blood, Tabor, Max-element, Tirant, Circe, gypsy10, GATE, mdg3, copia, Transpac, accord, 17.6, Stalker2, diver, gypsy5, gypsy6, McClintock, opus, gtwin, ZAM, mdg1, HMS-Beagle2, gypsy3
- non-LTR (3): non-LTR: Juan, G6, Het-A
- TIR (1): P-element

#### 3.8.3 burst of extant family in D. simulans

- LTR (1): diver2
- non-LTR (2): Cr1a, Rt1c
- TIR (6): hobo, transib2, 1136, Tc1, HB, (INE-1)

#### 3.8.4 burst of extant family in D. melanogaster

- LTR (9): 412, Burdock, flea, roo, HMS-Beagle, Quasimodo, 3S18, 297, Idefix
- non-LTR (8): Doc, jockey, R1A1-element, Ivk, Rt1b, Rt1a, F-element, X-element
- TIR (1): hopper;

### 3.9 Influence of the population size on equilibrium copy numbers

For computational reasons it was necessary to rescale the population size, which is assumed to be in the order of millions in *D. melanogaster* (Kreitman, 1983), to *N* = 10.000. However, as the standard deviation of equilibrium copy numbers decreases with increasing population size (e.g. using a sample of 400 simulations for an average copy number of 600: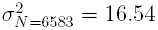, 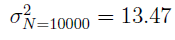, F-test *p* = 4.4*e* − 05) our approach is conservative.

